# SynThIA: A semi-automated tool for quantification of multi-partite synapses

**DOI:** 10.64898/2026.03.23.713591

**Authors:** Max Neather, James Morgan, Fong Kuan Wong

## Abstract

Synapses are evolutionarily conserved structures that form the fundamental units of neural communication. In the adult mouse cerebral cortex, most synapses are enveloped by glial protrusions from astrocytes and microglia, forming multi-partite synapses. Despite their prevalence, quantitative tools to systematically analyse these multi-cellular structures are limited to two or at most three markers. Here, we present Synapse Thresholding Image Analyser (SynThIA), an open-source, Python-based pipeline for high-throughput and accurate quantification of synapses, including multi-partite synapses. SynThIA enables multichannel analysis of up to four markers, providing detailed measurements of synaptic composition and distribution. The pipeline features an intuitive graphical interface allowing for users with minimal programming experience and a modular design that allows customization for advanced users. By combining accessibility and precision, SynThIA addresses a key methodological gap in multi-partite synaptic image analysis and provides a robust platform for studying synaptic organization in both *in situ* and *ex situ* preparation.

## INTRODUCTION

The synapse is a fundamental unit of neuronal communication, acting as a junction between neurons to facilitate information transfer that is essential for normal brain function. Understanding how synapses develop and operate is key to deciphering how the brain processes information and responds to its environment. Structurally, synapses are complex, multi-cellular compartments composed not only of neurons but also glial cells such as astrocytes and microglia^1^. From the neuronal perspective, the presynaptic terminal – located at the axonal end – transmits signals across the synaptic cleft through the release of neurotransmitters. These neurotransmitters bind to receptors on the postsynaptic membrane of the adjacent neuron, triggering electrical impulses known as action potentials. In addition to this neuronal framework, glial cells dynamically interact with synapses^1^. Astrocytes and microglial processes contact both presynaptic and postsynaptic compartments, though on different timescales^2,3^. Recent studies have shown between 30 to 60% of synapses in the adult mouse neocortex are contacted by glial processes at a given time^4^. These interactions regulate synaptic activity by modulating neurotransmitter release and receptor dynamics, playing essential roles not only in mature brain functions but also during neural circuit construction and neurodegeneration^5^.

To study synaptic behaviour, researchers have traditionally employed both *in situ* (e.g., brain slices) and *ex situ* (e.g., isolated synaptic terminals through synaptosome isolation) approaches^6^. Among *ex situ* techniques, synaptosome isolation offers a unique advantage – the preservation of isolated synaptic structure and molecular machinery, whilst enabling the investigation of synaptic transmission in a more controlled and accessible environment^7^. While synaptosome-based studies have primarily focused on neuronal components, these isolated synaptic structures can also retain associated glial elements^8–10^. This makes synaptosomes a valuable tool for studying multi-partite synapses.

Quantifying synaptic density and composition is central to assessing synaptic variation and dynamics. Commonly used tools such as Puncta Analyser and Synapse Counter – both FIJI plugins – enable quantification of pre- and postsynaptic markers in *in situ* conditions and can be adapted for *ex situ* applications^11,12^. However, these tools are limited to analysing only two fluorescent channels. Thus, it prevents the inclusion of glial processes in synaptic quantification. To address this limitation, a more recent tool, SynBot, was developed to quantify tripartite synapses involving glial, presynaptic and postsynaptic elements^13^. While SynBot’s three-channel capability marks a significant advancement, it does not account for other cellular interactions, nor does it provide detailed metrics such as counts of singly or doubly co-localized synapses (e.g., presynaptic and postsynaptic element colocalization or glial and synaptic element colocalization).

Here, we present Synapse Thresholding Image Analyser (SynThIA), an open-source, Python-based synapse quantification tool. SynThIA is a multi-channel analyser with a user-friendly graphical interface designed for high-throughput image processing. It enables detailed assessment of synaptic composition and can detect multi-partite synapses, including those involving both astrocytic and microglial components. Through customizable filtering options, SynThIA minimizes noise present in *ex situ* preparations. Notably, SynThIA is also compatible with *in situ* analysis. Finally, we have validated SynThIA by benchmarking its performance against SynBot and Synapse Counter. Collectively, we provide a robust, user-centric synapse analysis tool capable of batch processing and comprehensive multi-channel quantification.

## RESULTS

SynThIA is an open-source image analysis pipeline designed to analyse multi-channel synaptic images (Figure 1). We developed SynThIA in Python and integrated functions from the Scikit-image library for accessibility and flexibility^14^. The pipeline is user-friendly and requires no programming experience. However, its design allows more advanced users the opportunity to customize individual steps. All required files and installation instructions are freely available on Github (https://github.com/FKW-Lab/SynThIA/tree/main). We structured the pipeline around three main stages: image pre-processing, segmentation and analysis (Figure 1B).

**Figure 1.**
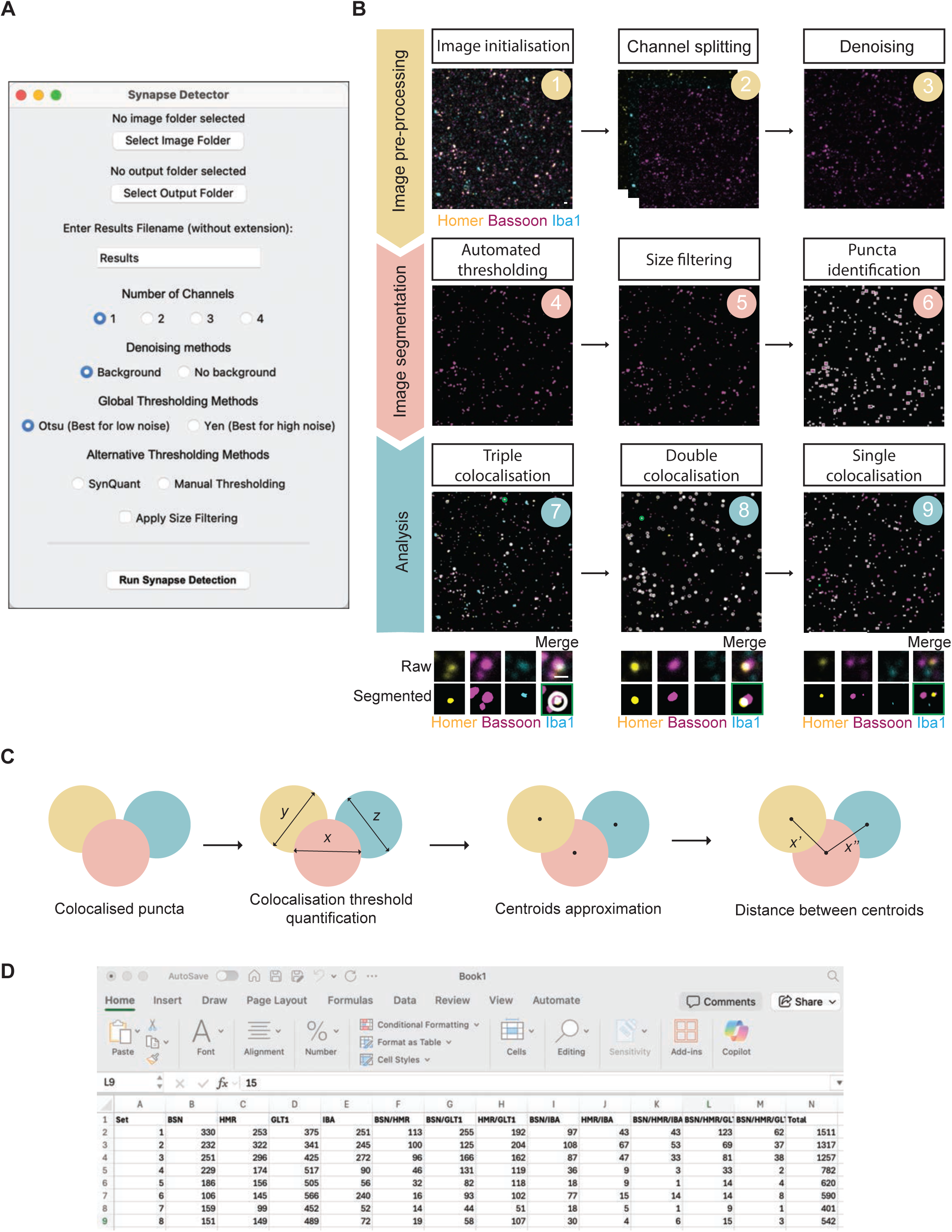
SynThIA image analysis pipeline and graphical user interface. (**A**) SynThIA graphical user interface opens automatically following user’s initialising Python. User can then load image(s) and set desired analysis settings. (**B**) Main steps in SynThIA analysis pipelines: (1) Image initialisation as user upload image(s), (2) Channels are assigned and split if loaded in .lif format, (3) Images are denoised, (4) User selected threshold is applied, (5) Size filter applied if selected by user, (6) Connected groups of pixels labelled as individual puncta, (7) Triple colocalization events quantified, (8) Double colocalization events quantified, (9) Remaining puncta not colocalised is quantified as single puncta. Green circles in Figure 7-9 are selected as examples of triple- and double-colocalised and single puncta enlarged below, respectively. (**C**) Circulation approximation for colocalization quantification. SynThIA quantifies the colocalization threshold based on the average diameter of the puncta from the different channels (yellow, red, and blue circles). Next, it identified the centroid within each signal and measured the distance between centroids. Any puncta that fall below the colocalization threshold are identified as colocalised (Note: The distance between puncta is quantified and compared to the colocalization threshold in the master channel (e.g., red channel)). (**D**) Example of excel output from SynThIA Scale bar: 1µm

### Preparing and pre-processing images

As with any image analysis workflow, the reliability and accuracy of the output are inherently dependent on the image quality. We therefore recommend that users prioritize the acquisition of high-quality images with consistent, high signal-to-noise ratios. Here, we typically use a confocal microscope and image our samples with 1024 x 1024 resolution. The first stage of image analysis in SynThIA involves image preparation. Users begin by specifying the location of their datasets, which can be supplied as single multichannel images (image set) or as by batch files such as *.lif* datasets (Figure 1A). For single image set, channels must be manually separated and named following the naming convention x.y where x denotes the image set and y denotes the specific channel (e.g., 1.2 for channel 2 of a set of named ‘1’). Batch files are automatically split into individual channels by SynThIA (Figure 1B, Step 2). Once defined, the output directory will store both quantification results and a series of validation images that document each processing step. These validation files are particularly important for assessing whether the chosen parameters are appropriate and can guide further refinement when necessary. SynThIA supports the processing of up to four image channels.

Pre-processing begins by splitting the images into individual channels and converting each channel to grayscale (Figure 1B). To reduce background signals and enhance the visibility of synaptic puncta, we have included a denoising step (Figure 1B, Step 3). To account for the different levels of noise inherent in different sample types and preparation, SynThIA offers two denoising strategies. For images that tend to exhibit high noise, such as those from tissues, a Difference of Gaussian filter is applied as it enhances the puncta edge^15^. This is followed by a rolling ball filter that reduces the background noise^16^. For images with low levels of noise, such as those from synaptosome isolation, similar to other image analysis pipelines, SynThIA instead applies an Anscombe transform to stabilize Poissonian noise, followed by a 3x3 median filter and non-local means denoising^17,18^. All denoising parameters can be tuned on a channel-specific basis with minimal coding required to make these adjustments. We have provided annotations in our code for users to make the appropriate adjustments accordingly.

### Detecting and segmenting synaptic puncta

The second stage focusses on segmentation, the process of distinguishing puncta from the background. Due to the size and structure of synaptic puncta, these puncta often resemble noise, especially in images obtained from isolated synaptosome. To circumvent this, we have included two key steps for image segmentation: thresholding and size filtering (Figure 1B, steps 4 and 5). Thresholding classifies each pixel as either foreground or background. As synaptic puncta can vary in intensity and shape, the choice of thresholding method can strongly influence the results. SynThIA provides multiple options, including global thresholding methods such as Otsu, which is generally less aggressive and suitable for many image types and Yen, which is more effective for highly noisy datasets. Alternatives such as manual thresholding and SynQuant are also available (Figure 1A)^19^. Importantly, SynThIA’s modularity allows users to add other thresholding functions from the Scikit-Image library with minimal effort into the provided code, consequently adding to the versatility and usability of the pipeline. After image thresholding, each channel is labelled using the Skimage label function, where any connected pixels with greyscale values above 0 are treated as a single punctum (Figure 1B, step 6).

We have included the option to further refine puncta identification using size filtering (Figure 1B, step 5). Size filters are defined by the user based on the expected size of the puncta. This can be estimated by using tools such as Fiji’s ‘Analyze Particles’ function. If a size filter has been selected, the areas of the labelled puncta are calculated using the Skimage regionprops function and compared against these boundaries. Any object falling outside the expected size range are exclude from the dataset, ensuring that only relevant puncta are retained further downstream analysis.

### Quantitative analysis

In the final stage, SynThIA identifies colocalised and non-colocalised puncta across the channels. First, puncta diameters are estimated from the areas, assuming circular geometry by applying the formula,

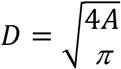

A channel-specific mean diameter is then calculated and used to define a colocalization threshold distance (Figure 1C). For multi-channel datasets, the threshold is set to be the average diameter across channels analysed (i.e., In an image set containing three channels A, B, and C, the double colocalization threshold for channels A and B will be different for channels A and C). Centroids of puncta are identified using the Skimage regionprops function. In SynThIA, colocalization is determined relative to a designated master channel, which is set by default to channel 1. For the master channel punctum, the nearest centroids in the other channels are identified and their distances compared to the threshold (Figure 1C). If all distances fall below the threshold, the puncta are considered colocalised. Colocalised puncta are then ignored in subsequent analysis to prevent double-counting. Colocalization analysis proceeds hierarchically, first identifying quadruple-colocalised puncta (when applicable), followed by triple- and double-colocalised puncta (Figure 1B, steps 7 and 8). Any remaining puncta that are unpaired are identified as single puncta. For batch datasets, this process is iterated in a series of loops across all images until the analysis is complete. Once completed, the analysed dataset is exported into an excel file, providing detailed metrics of the composition of both single and colocalised events (Figure 1D).

### Validation of SynThIA using two-channel simulated synaptosome images

To evaluate the accuracy and robustness of SynThIA, we first generated simulated synaptic images consisting of two 10 x 10 grids of puncta derived from images of previous synaptosome datasets (Figure S1A). In these simulations, when channels were merged, all puncta are double-colocalized. To test SynThIA’s ability to correctly detect these events and discriminate puncta from noise, we applied varying levels of random intensity Gaussian noise to the respective percentage of unoccupied pixels (2.5% to 50%) in the individual images (Figure S1B). These noise levels were selected to reflect those commonly encountered in typical experimental datasets (10 to 30%) as well as more extreme conditions (50%). Three independent sets of noise-simulated images were generated using randomised filters to avoid bias from uneven pixel distributions.

Precision was calculated as the number of correctly identified colocalization events (true positives) divided by the total number of detected events (e.g., true positives and false positives). Using global thresholding methods such as Otsu and Yen, both approaches achieved high precision up to 20% noise, with no significant differences observed between methods (Figure S1C). At higher noise levels, precision declined for both, reflecting an increase in false positive detection. Incorporating size filtering after thresholding substantially mitigated this decline, maintaining high precision across all noise levels (Figure S1C).

We next compared SynThIA’s performance with other established synapse quantification pipelines, including Synapse Counter and SynBot (with ilastik thresholding; Figures S1D and S1E)^12,13,20^. For both SynThIA and Synapse Counter, size filtering significantly improved precision (Figure S1D) and accuracy (i.e., correct discrimination between puncta and noise; Figures S1D and S2). Without size filtering, both pipelines exhibited elevated false positives where noise was erroneously classified as single puncta events (Figures S1E and S2). In contrast, SynBot performance was unaffected by the addition of size filtering. Its baseline precision remained consistently lower than SynThIA across all noise conditions. (Figure S1D).

To further assess SynThIA’s capacity to distinguish colocalised puncta from single events, we randomly deleted 30 of the 100 puncta from each channel in the simulated images (Figure 2A). This resulted in datasets containing 49 colocalised puncta and 21 single puncta per channel. As before, random Gaussian noise was applied to the respective percentage of unoccupied pixels at levels ranging from 2.5% to 50% (Figure 2B). Across most noise levels observed in typical experimental datasets, SynThIA with size filtering outperformed both Synapse Counter and SynBot in detecting both single and double events (Figures 2C and 2D). At the highest noise level (50%), SynThIA’s performance decreased markedly, failing to reliably distinguish between puncta from background noise. By contrast, SynBot maintained relatively stable performance up to 50% noise. Although its absolute precision decreased by ∼23% at 50% noise compared to no noise (Figure 2C), it remained effective in distinguishing puncta from background noise (Figure 2D). Overall, these benchmarking experiments demonstrate that SynThIA provides reliable and accurate quantification of single and double synaptic events under realistic noise conditions and outperformed both Synapse Counter and SynBot. However, under extreme noise levels outside of typical experimental noise, SynBot outperformed both SynThIA and Synapse Counter especially in its capacity to distinguish single events from noise.

**Figure 2.**
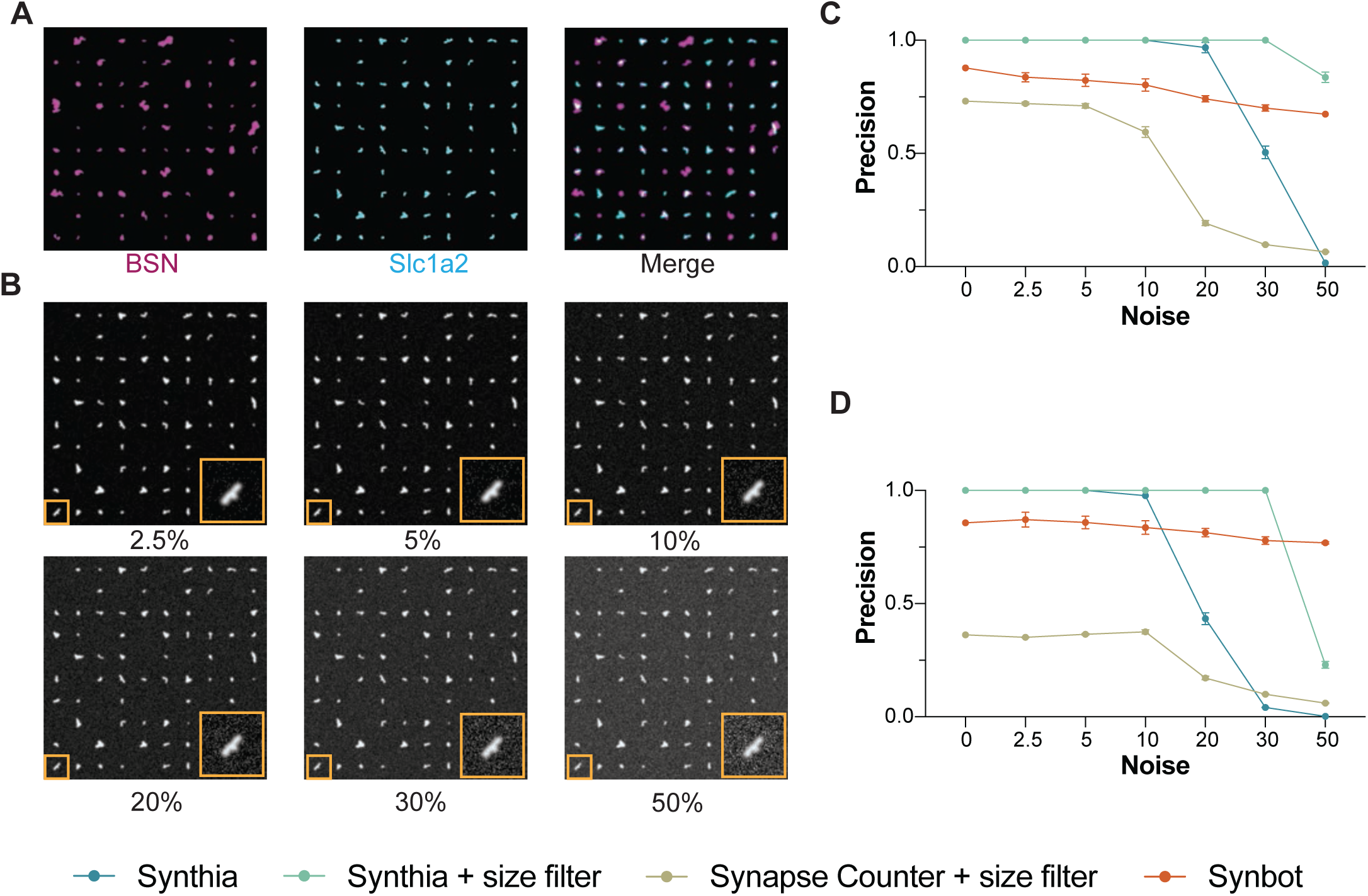
SynThIA validation on simulated synapse images with different noise levels. (**A)** Simulated synapse images generated from prior synaptosome dataset stained with Bassoon (magenta) and Slc1a2 (Cyan) arranged in 10 x 10 grids of puncta. In these images, we have randomly deleted 30 of the 100 puncta from each channel. (**B**) Simulated synapse images from prior synaptosome dataset stained with Slc1a2 (white) applied with varying levels of random intensity of Gaussian noise (2.5% to 50%) to the unoccupied pixels in the image. Yellow insets show selected simulated puncta in higher magnification. (**C**) Precision (true positives/total detected events) measurement obtained using SynThIA (Otsu, cyan), SynThIA with size filtering (Otsu, green), Synapse counter with size filtering (olive green), and SynBot (orange) for double-colocalised events. 2-way ANOVA (*F_interaction_* (18,56) = 198.7, *p* <0.0001). Three independent sets of noise-simulated images were generated at each noise level. Results are expressed as mean ± standard error of mean (SEM). (**D**) Precision (true positives/total detected events) measurement obtained using SynThIA (Otsu, cyan), SynThIA with size filtering (Otsu, green), Synapse counter with size filtering (olive green), and SynBot (orange) for single events. 2-way ANOVA (*F_interaction_* (18,56) = 285.9, *p* <0.0001). Three independent sets of noise-simulated images were generated at each noise level. Results are expressed as mean ± standard error of mean (SEM).

### Validation of SynThIA using synaptosome images

We next evaluated SynThIA using experimental synaptosome images obtained from adult mouse cortical preparations stained with presynaptic (Bassoon+), postsynaptic (Homer+), and perisynaptic astrocytic protrusion (PAP, Slc1a2+ also known as Excitatory Amino Acid Transporter 2 (Eaat2)) markers (Figures 3 and S3). Similar to previous reports, we consistently observe a higher proportion of presynaptic than postsynaptic events in our synaptic isolations^21^. In addition, ∼38% of cortical synaptic puncta colocalised with PAP (i.e., Bassoon+ Slc1a2+, Homer+ Slc1a2+ and Bassoon+ Homer+ Slc1a2+), aligning with values previously reported in adult mouse cerebral cortex (∼30 - 60%)^4^. A key feature of SynThIA is its ability to quantify both single and colocalised events, providing a comprehensive overview of the synaptic distributions in the dataset (Figures 3C and 3D). Most puncta represented single synaptic events, with ∼31% Bassoon+ and ∼16% Homer+ puncta. In addition, ∼17% of puncta were double-labelled for Bassoon and Homer, and ∼8% of puncta were tripled-labelled (Bassoon+, Homer+, Slc1a2+) (Figure 3D).

**Figure 3.**
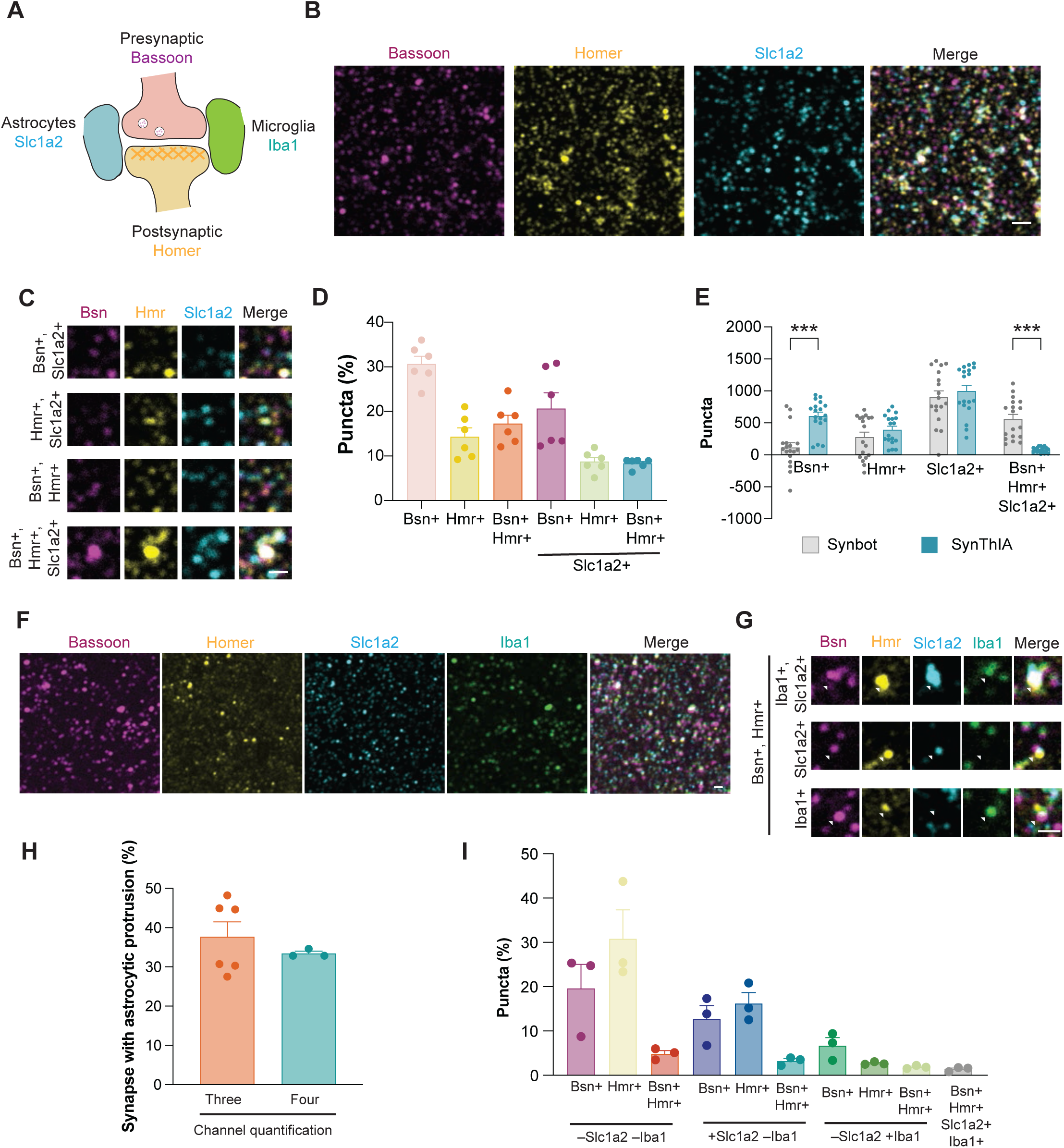
SynThIA quantifies multi-partite synapses from isolated synaptosomes. (**A**) Schematic of multi-partite synapses (**B**) Cortical synaptosomes isolation from Sighted-FVB mice at P56 following immunohistochemistry for Bassoon (magenta), Homer (yellow), and Slc1a2 (cyan). (**C**) Higher magnification of the cortical synaptosomes isolated from Sighted-FVB mice at P56. (**D**) Quantification of percentage of synaptic puncta from synaptosomes isolated from Sighted FVB mice at P56 that are Bassoon+ (pink), Homer+ (yellow), Bassoon+ Homer+ (orange), Bassoon+ Slc1a2+ (Magenta), Homer+ Slc1a2+ (green), and triple colocalised (cyan). One-Way ANOVA (*F* (5,30) = 17.31). N = 6 mice (**E**) Quantification of synaptic puncta from synaptosomes isolated from Sighted FVB mice at P56 quantified using SynBot (grey) and SynThIA (cyan) for Basson+, Homer+, Slc1a2+, and triple colocalised. Two-Way ANOVA with Sidak multiple comparison (*F_interaction_* (3,136) = 15.61, *** *p* <0.001). N = 18 wells from 6 mice. **(F)** Cortical synaptosomes isolation from Sighted-FVB mice at P56 following immunohistochemistry for Bassoon (magenta), Homer (yellow), Slc1a2 (cyan) and Iba1 (green). (**G**) Higher magnification of the synaptosomes isolated from Sighted-FVB mice at P56. (**H**) Quantification of percentage of synaptic puncta with astrocytic processes from the three-channels (orange) and four-channels (green) quantifications. Mann-Whitney test. N = 6 mice (Three channels), 3 mice (Four Channels). (**I**) Quantification of the percentage of synaptic puncta from synaptosome isolated from Sighted FVB mice at P56 that are Bassoon+ (magenta), Homer+ (yellow), Bassoon+ Homer+ (orange), Bassoon+ Slc1a2+ (dark blue), Homer+ Slc1a2+ (light blue), Bassoon+ Homer+Slc1a2+ (cyan), Bassoon+ Iba1+ (Dark green), Homer+ Iba1+ (green), Bassoon+ Homer+ Iba1+ (light green), and quadruple colocalised (grey). One-Way ANOVA (*F* (9,20) = 10.32). N = 3 mice. Results are expressed as mean ± standard error of mean (SEM). The data points within the bar graphs indicate the average percentage per animal (D), (H), and (I) average puncta per well (E). Scale bars: 1µm.

To benchmark SynThIA against existing tools, we compared it to SynBot (Figures 3 and S3)^13,20^. We were unable to compare SynThIA to Synapse Counter as its analysis is limited to only two channels^12^. Unlike SynThIA, SynBot does not provide a full breakdown of the synaptic distribution of the dataset^13^. Instead, it reports total single puncta regardless of colocalization, along with the total number of triple colocalization events (Figure S3D). For this reason, we compared total puncta and triple colocalization counts between pipelines. SynBot consistently detected higher event counts across conditions, except for Bassoon+ puncta (Figure S3D). Closer inspection revealed several factors contributing to this discrepancy: 1) SynBot combined with ilastik thresholding was more sensitive to low-intensity fluorescent signals (Figures S3A and S3B), 2) segmented puncta in SynBot often exhibited enlarged areas (Figures S3A and S3B), and 3) individual colocalised puncta were counted multiple times (Figures S3B and S3C). This double-counting was evident in 36% of the image set quantified, where the total number of puncta in a single channel was lower than the number of triple-colocalised puncta identified (Figure 3E). In contrast, SynThIA was specifically designed to prevent double-counting, as once a punctum is classified as colocalised (double or triple), it is removed from the pool of events. This ensures a non-redundant synaptic quantification.

To demonstrate SynThIA’s capacity for multi-channel analysis beyond three markers, we stained synaptosomes for Bassoon, Homer, Slc1a2, and the microglial marker, Iba1 (Figures 3A, F, G). The fraction of synapses associated with astrocytic processes quantified by SynThIA was comparable between the three-(∼37%) and four-channel (∼33%) analyses (Figures 3H). Furthermore, in the four channel quantifications, ∼11% of synapses colocalised with microglial protrusions and ∼1.4% with both astrocytic and microglial elements (Figures 3H, I). Thus, SynThIA enables detailed quantification of synaptic composition across multiple cell types including the ‘quad-partite’ synapses. Collectively, these analyses demonstrate that SynThIA provides more granular and accurate quantification of synaptic events by distinguishing uncolocalised, double- and triple-colocalization, thereby offering a more faithful representation of synaptic organisation from our dataset.

### Application of SynThIA across diverse synaptic markers

Bassoon and Homer are well established scaffolding proteins localised at presynaptic and postsynaptic sites, respectively^22,23^. To assess whether SynThIA can reliably identify additional synaptic proteins, we next examined synaptic markers located in distinct subcellular compartments. Specifically, we assessed Snap25, a plasma membrane-anchored SNARE protein involved in synaptic vesicle function, and Grin2B (also known as GluN2B), a postsynaptic NMDA receptor subunit (Figure 4A)^24,25^. Among the identified synaptic puncta, ∼51% of puncta were colocalised, while ∼30% and ∼20% were positive for Snap25 and Grin2b alone, respectively (Figures 4B and E).

**Figure 4.**
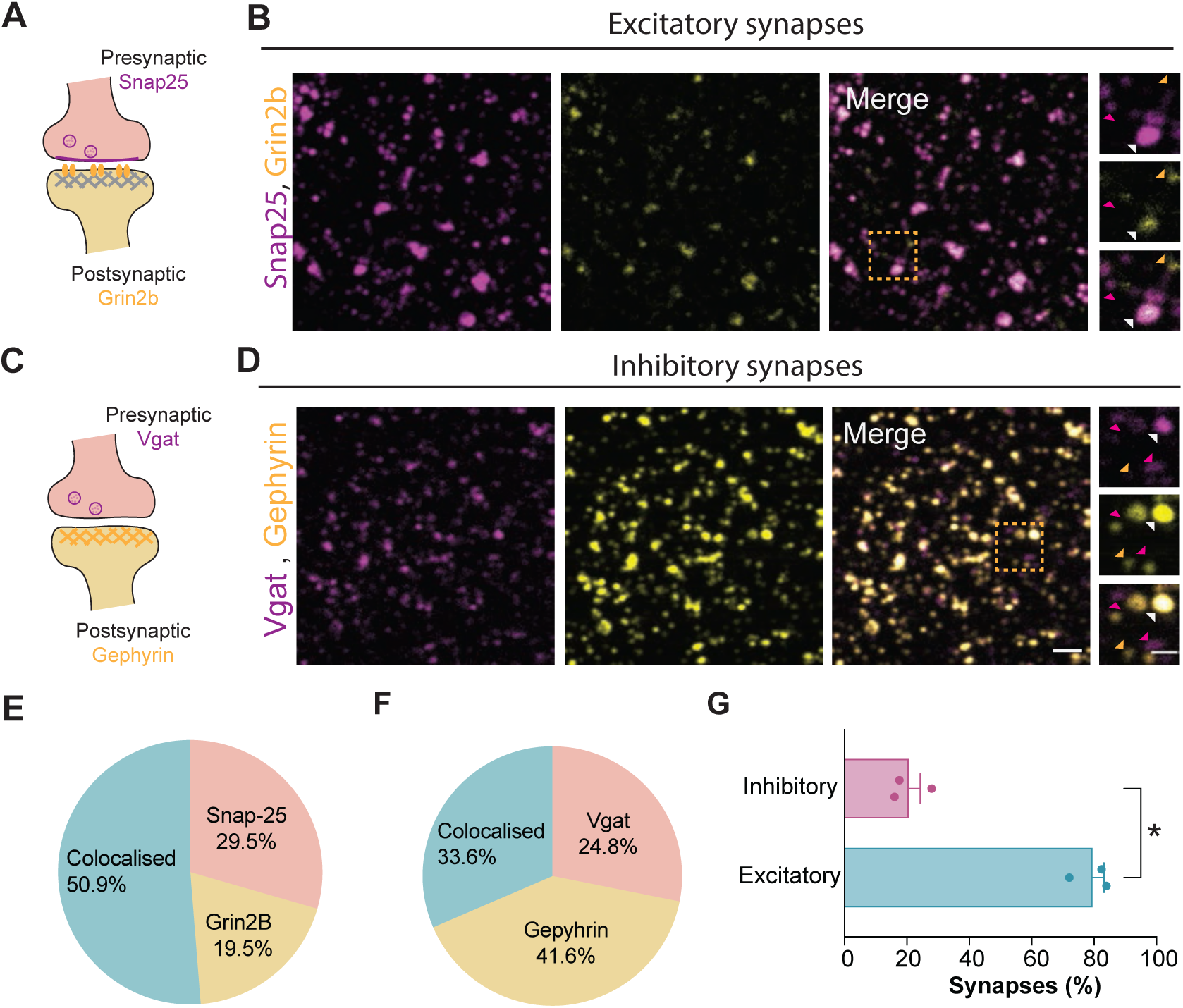
SynThIA quantifies excitatory and inhibitory synapses from isolated synaptosomes. (**A)** Schematic of an excitatory synapse with Snap25 and Grin2b localisation. (**B**) Cortical synaptosomes isolation from Sighted-FVB mice at P60 following immunohistochemistry for Snap25 (magenta) and Grin2b (yellow). Dashed yellow box shows the amplified area in the images to the right. Magenta arrowheads exemplified Snap25+ Grin2b-, yellow arrowhead for Snap25-Grin2b+, and white arrowhead for Snap25+ Grin2b+ synaptic puncta. (**C**) Schematic of an inhibitory synapse with Vgat and Gephyrin localisation. (**D**) Cortical synaptosomes isolation from Sighted-FVB mice at P60 following immunohistochemistry for Vgat (magenta) and Gephyrin (yellow). Dashed yellow box shows the amplified area in the images to the right. Magenta arrowheads exemplified Vgat+ Gephyrin-, yellow arrowhead for Vgat-Gephyrin+, and white arrowhead for Vgat+ Gephyrin+ synaptic puncta. (**E**) Quantification of percentage of synaptic puncta from synaptosome isolated from Sighted FVB mice at P60 that are Snap25+ (red), Grin2b+ (yellow), Snap25+ Grin2b+ (cyan). N =3 mice. (**F**) Quantification of percentage of synaptic puncta from synaptosome isolated from Sighted FVB mice at P60 that are Vgat+ (red), Gephyrin+ (yellow), Vgat+ Gephyrin+ (cyan). N = 3 mice. (**G**) Quantification of the percentage of excitatory (cyan) and inhibitory (magenta) synapses at P60. Paired t-test, \**p* = 0.0157, N = 3 mice. Results are expressed as mean ± standard error of mean (SEM). The data points within the bar graphs indicate the average per animal. Scale bars: 2µm, 1µm (inset).

We then investigated whether SynThIA could also quantify inhibitory synapses in our synaptosome preparations. To this end, we stained for Slc32a1 (also known as vesicular γ amino acid (Vgat)), a presynaptic vesicular GABA transporter, together with Gephyrin, a postsynaptic scaffolding protein critical for inhibitory synapse assembly (Figure 4C)^26,27^. Approximately 33.6% of puncta showed Vgat+ Gephyrin+ colocalization, while 24.8% and 41.6% were positive for Vgat and Gephyrin alone, respectively (Figures 4D and 4F).

Previous studies have shown that ratio of excitatory to inhibitory synapses varies across brain areas and cell types. In most brain regions, excitatory synapses greatly outnumber inhibitory synapses, with reported ratios ranging from 3:1 to 7:1^28,29^. In our synaptosome isolations, when considering only double-colocalised excitatory (Snap25+ Grin2b+) and inhibitory (Vgat+ Gephyrin+) synapses in the same samples, we observed a ratio of approximately 5:1 (Figure 4G). This provides further validation of the accuracy of SynThIA quantification. Together, these findings demonstrate that SynThIA can quantify synaptic proteins across distinct subcellular compartments and can be applied to both excitatory and inhibitory synapses.

### Validation of SynThIA in *in vivo* excitatory synapses

SynThIA was developed to quantify synapses in both *in situ* and *ex situ* approaches. While all preceding experiments were validated using *ex situ* approaches (synaptosome isolation), we next assessed SynThIA’s ability to quantify synapses directly from mouse brain tissue sections. To this end, we analysed excitatory synapses colocalised with astrocyte protrusions by staining for Bassoon, Homer and Slc1a2, in the CA1 hippocampus of wild-type adult mice and in *Fragile X messenger Ribonucleoprotein1* knockout (*Fmr1-/-*) mice (Figure 5A). Previous work using electron microscopy have reported an approximately 10% reduction in synapses with astrocytes in the CA1 region of *Fmr1-/-* mice^30^. Using SynThIA, we observed a comparable ∼15% decrease in the percentage of synapses associated with astrocytic protrusions in *Fmr1-/-* mutant mice in the CA1 region of the hippocampus(Figures 5B-D). Together, these findings validate the accuracy of SynThIA in quantifying synapses in both *in situ* and *ex situ* settings. Notably, the *in situ* quantifications obtained using SynThIA closely matched values derived from electron microscopy – the current gold standard for synaptic analysis.

**Figure 5.**
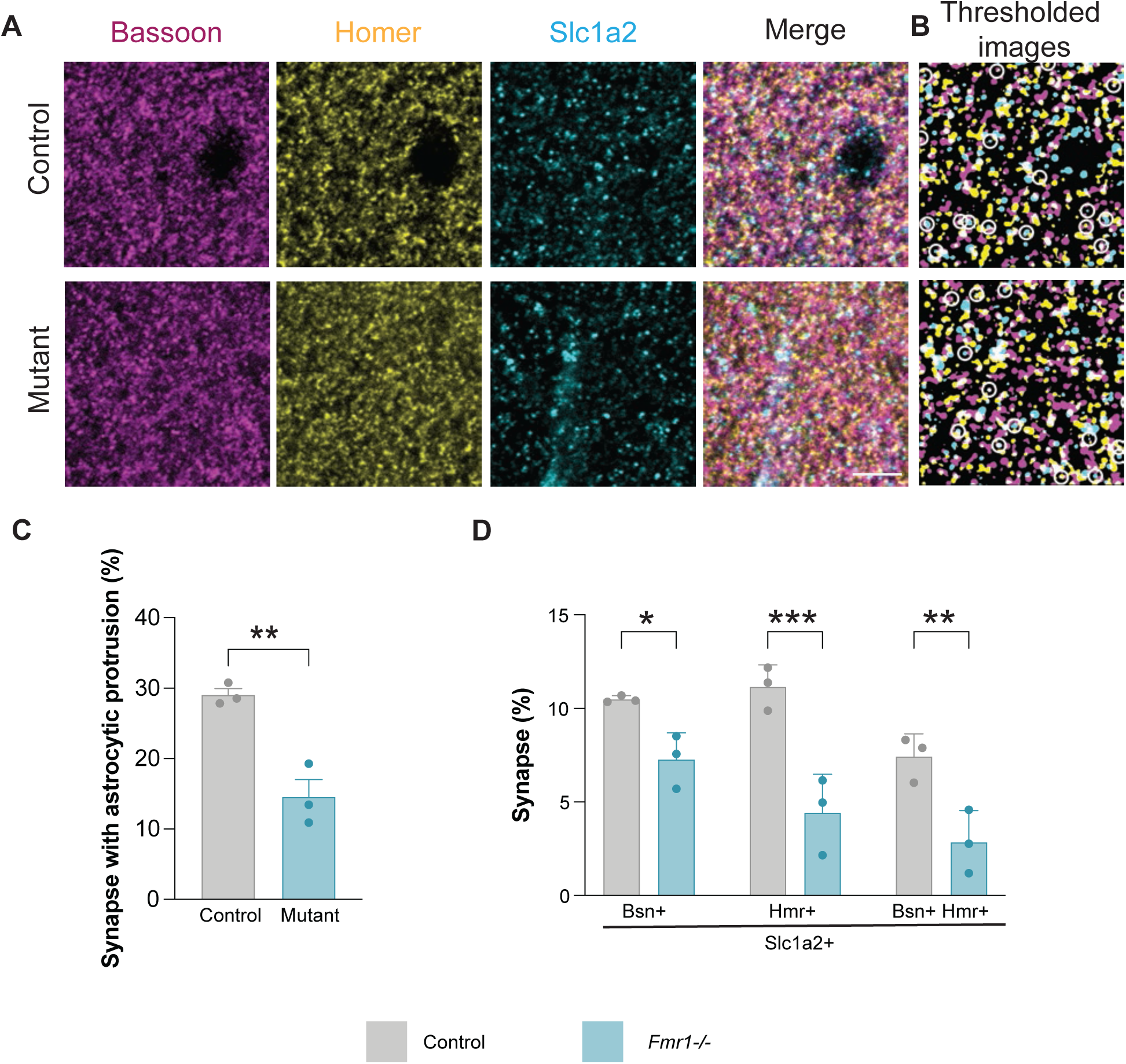
SynThIA quantifies excitatory synapses *in vivo* accurately and reliably. (**A**) Coronal sections of CA1 of the hippocampus of Sighted-FVB (control, top) and *Fmr1-/-*(bottom) at P60 following immunohistochemistry for Bassoon (magenta), Homer (yellow), and Slc1a2 (cyan). (**B**) Output images identifying triple colocalization (white circle) produced by SynThIA in control (top) and *Fmr1-/-* (bottom). (**B**) Quantification of the percentage of synapses with astrocytic protrusion in control (grey) and *Fmr1-/-* (cyan). Two-tailed Student’s unpaired t-test, \*\**p* = 0.0052, N = 3 mice. (**C**) Quantification of percentage of Bassoon+, Homer+, and Bassoon+ Homer+ with astrocytic protrusion in control (grey) and *Fmr1-/-* (Cyan). Two-way ANOVA (*F_interaction_* (2, 12) = 2.233). \**p* = 0.049, \*\**p* = 0.0057, \*\*\**p* = 0.0002. N=3 mice. Results are expressed as mean ± standard error of mean (SEM). The data points within the bar graphs indicate the average per animal. Scale bar: 5µm,

## DISCUSSION

Despite the growing recognition of glial involvement in synaptic transmission^5^, methodological limitations have constrained the systematic characterization of these complex, multicellular synaptic structures. There remains a lack of accessible analytical tools capable of reliably and accurately quantifying multicellular synaptic compartments across multiple fluorescent channels in a high-throughput manner. To address this methodological gap, we developed SynThIA, a multichannel synaptic analysis pipeline designed for accurate and efficient quantification of up to four channels. SynThIA supports both *in situ* and *ex situ* preparations, generating high-resolution granular data on synaptic composition and distribution. This enables detailed interrogation of physiological and/or pathological synaptic changes in any neurological systems. Importantly, SynThIA is an open-source platform accommodating a broad spectrum of users: it integrates an intuitive graphical user interface for researchers with little to no computational experience, while its modular architecture allows customization for advanced users.

SynThIA was designed to overcome common challenges in image analysis, such as variations in signal-to-noise ratios through its integrated preprocessing and analysis pipeline. Recognising that the complete elimination of noise is rarely feasible in any experimental settings, SynThIA incorporates two distinct denoising strategies optimized for low- and high-noise samples – an option rarely available in comparable analysis tools (Table S2). In both strategies, emphasis was placed on preserving and enhancing puncta edges^15,17^. As a result, the shapes of identified puncta remain well preserved and not artificially enlarged, providing a faithful representation of the original signals essential for subsequent quantitative analyses (Figure S3). However, there are inherent trade-offs in the current denoising and thresholding strategies. For instance, in high-noise images, the Difference of Gaussian approach proved most effective for enhancing puncta boundaries. Yet, as this method relies on the subtraction of two Gaussian-blurred versions of the image with different standard deviations, puncta with weaker intensity values and/or diffused edges are often attenuated or lost during processing^15^. This effect was evident in our benchmarking results, where SynThIA, compared to SynBot, occasionally omitted low-intensity puncta and showed a mild bias toward higher-intensity structures. In contrast, SynBot coupled with ilastik based thresholding extracts features primarily based on puncta shape; as a result, its sensitivity depends strongly on user-provided training data and quality and diversity of images used for training (e.g., the range of puncta shapes and sizes).

SynThIA was also developed with high-throughput and rapid processing as a key design principle. In addition to incorporating batch processing capabilities, several strategies were implemented to reduce computational demands and processing time (Table 2). This includes the use of global thresholding algorithms such as Otsu and Yen, which require no prior training data, thereby minimizing user bias and maintaining consistent segmentation performance. Furthermore, SynThIA employs a circular approximation approach for signal colocalization. By comparing the distance between centroids to the colocalization threshold, this reduces the computational burden and processing time to determine colocalization as opposed to scanning for individual pixel/voxel in the pixel overlap approach^31^. Thus, striking a balance between computational efficiency and spatial accuracy. These design choices substantially reduce runtime where in our benchmarking, processing 72 image sets required over 1.5 hours with SynBot, but only 30 minutes using SynThIA (Table 2).

When comparing SynThIA with existing synapse quantifying tools, SynThIA demonstrated clear advantages in both simulated and experimental datasets. Synapse Counter improved in precision when paired with size filtering but is limited by its two-channel capacity, restricting its utility for the identification of more complex synaptic architectures. Furthermore, its accuracy and precision in identifying single– and colocalised puncta was lower than that achieved by SynThIA (Figures S1 and 2). SynBot, particularly when used with ilastik-based thresholding, maintained stable performance under high-noise conditions and showed greater sensitivity to low-intensity signals. However, this increased sensitivity was associated with inflated puncta areas, elevated false positives, and frequent double-counting of colocalised events. While these limitations may be reduced through additional training, this requires significant user time and optimisation. Nonetheless, for images with typical experimental noise, SynThIA outperformed SynBot. In contrast, SynThIA was specifically designed to minimize processing time, prevent redundant event assignment and provide a structured breakdown of single, double, triple and quadruple colocalizations across channels, while maintaining high precision especially under typical noise conditions. This enabled more accurate quantification of synaptic organisation, including the ability to profile multi-cellular interactions such as astrocytic and microglial associations. Together, these comparisons highlight that while SynBot offers robustness under extreme noise especially in distinguishing signal from noise and Synapse Counter provide a rapid, descriptive information albeit with limited accuracy and channel numbers, SynThIA delivers better performance under typical noise conditions. Furthermore, it offers a more granular and biologically informative representation of synaptic distributions, particularly in multi-channel datasets.

Looking ahead, SynThIA’s application is not restricted to synaptic imaging. The pipeline can be readily adapted to other imaging modalities that require multichannel quantification of punctate or spot-like structures, such as single-molecule fluorescence in situ hybridization (smFISH). Its open-source and modular design encourage community-driven extension and adaptation. Further development could incorporate probabilistic segmentation models, such as those implemented in SynQuant, to further address background-related challenges^19^. However, such models increase computational complexity and may limit generalizability if overly tailored to specific markers. Alternatively, user-trainable classifiers such as ilastik offer dynamic, on-the-fly model generation, but this flexibility comes at the cost of increased computational intensity and the need for representative training datasets^20^.

### Limitations of the study

To reduce computational burden and processing time, SynThIA employs a circular approximation with automated thresholding to determine signal colocalization. Unlike pixel-overlap approaches, which assess direct voxel or pixel coincidence, the circular approximation evaluates spatial proximity, where two objects are considered colocalized only if they fall within a defined colocalization threshold. This threshold determines the degree of spatial association required for colocalization. Currently, SynThIA automatically sets this threshold as the average diameter of puncta across all channels. However, if the punctum in a specific channel is heterogeneous (e.g., differing in size and shape), this may affect colocalization accuracy. Therefore, users are encouraged to visually inspect output images to verify the suitability of the applied colocalization parameters for their specific dataset. A further limitation of the current implementation is that SynThIA is designed two-dimensional images, including both optical sections and z-stack projections. Unlike advanced commercial software such as Imaris, SynThIA does not support three-dimensional synaptic quantification. Finally, SynThIA was developed from a neuron-centric perspective, aimed at investigating the composition and distribution of multicellular synapses. While effective in quantifying the proportion of synapses containing glial processes, previous studies have shown that a single synapse can be contacted by two astrocytes^32–34^. Due to the hierarchical quantification approach which removes puncta once quantified, SynThIA cannot currently identify neuronal synapses contacted simultaneously by two glial protrusions from the same marker.

## SUPPLEMENTAL INFORMATION

## AUTHOR CONTRIBUTIONS

M.N., J.M., and F.K.W. conceived the study. M.N. performed most experiments described in this manuscript. J.M. contributed to data collection. M.N. and F.K.W. wrote the paper with inputs from all authors.

## ACKNOWLEDGEMENTS

We thank Takashi Namba, Emre Düşünell, Cerys Manning and Clémence Bernard for critical reading of the manuscript, and members of the Wong laboratories for stimulating discussions and ideas. This work was supported by grants from the Medical Research Council MR/T030143/1 (F.K.W.) and University of Manchester (F.K.W.).

---

**Figure S1.**
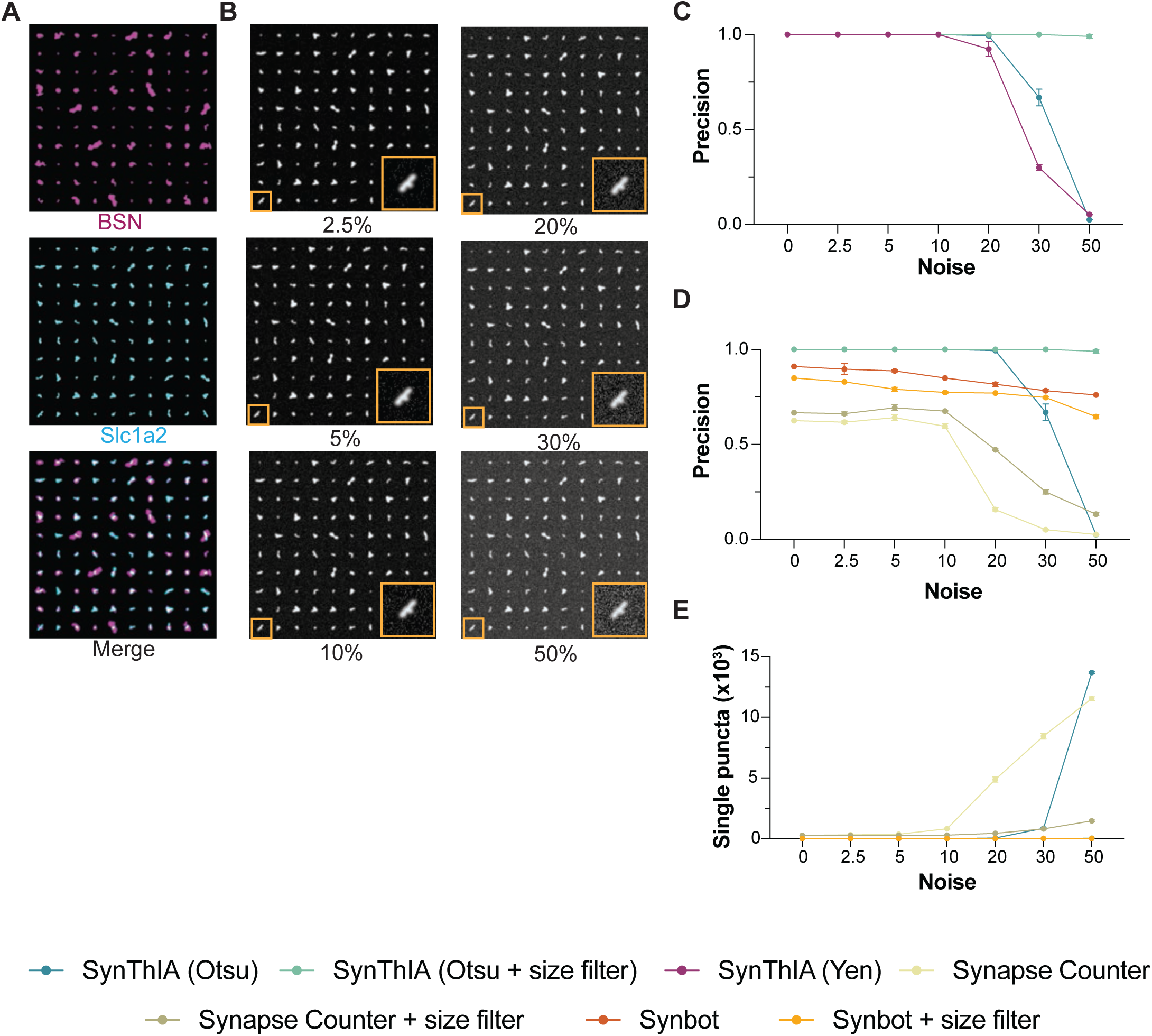
SynThIA validation on simulated synapse images with different noise level related to Figure 2. (**A**) Simulated synapse images generated from prior synaptosomes dataset stained with Bassoon (magenta) and Slc1a2 (cyan) arranged in 10 x 10 grids of puncta. (**B**) Simulated synapse images generated from prior synaptosome dataset stained with Slc1a2 (White) applied with varying levels of random intensity of Gaussian noise (2.5% to 50%) to the unoccupied pixels in the image. Yellow insets show selected simulated puncta in higher magnification. (**C**) Precision (true positives/total detected events) measurement obtained using SynThIA (Otsu, cyan), SynThIA with size filtering (Otsu, green), and SynThIA (Yen, Magenta) for double-colocalised events. 2-way ANOVA (*F_interaction_* (12, 42) =865.9, *p* <0.0001). Three independent sets of noise-simulated images were generated at each noise level. (**D**) Precision (true positives/total detected events) measurement obtained using SynThIA (Otsu, cyan), SynThIA (Otsu) with size filtering (green), Synapse counter (light green), Synapse counter with size filtering (olive green), SynBot (Orange), and SynBot with size filtering (yellow) for single events. 2-way ANOVA (*F_interaction_* (30, 84) = 728.7, *p* <0.0001). Three independent sets of noise-simulated images were generated at each noise level. (**E**) Quantification of single puncta detected using SynThIA (Otsu, cyan), SynThIA (Otsu) with size filtering (green), Synapse counter (light green), Synapse counter with size filtering (olive green), SynBot (Orange), and SynBot with size filtering (yellow) for single events. 2-way ANOVA (*F_interaction_* (30, 84) = 5657, *p* <0.0001). Three independent sets of noise-simulated images were generated at each noise level. Results are expressed as mean ± standard error of mean (SEM).

**Figure S2.**
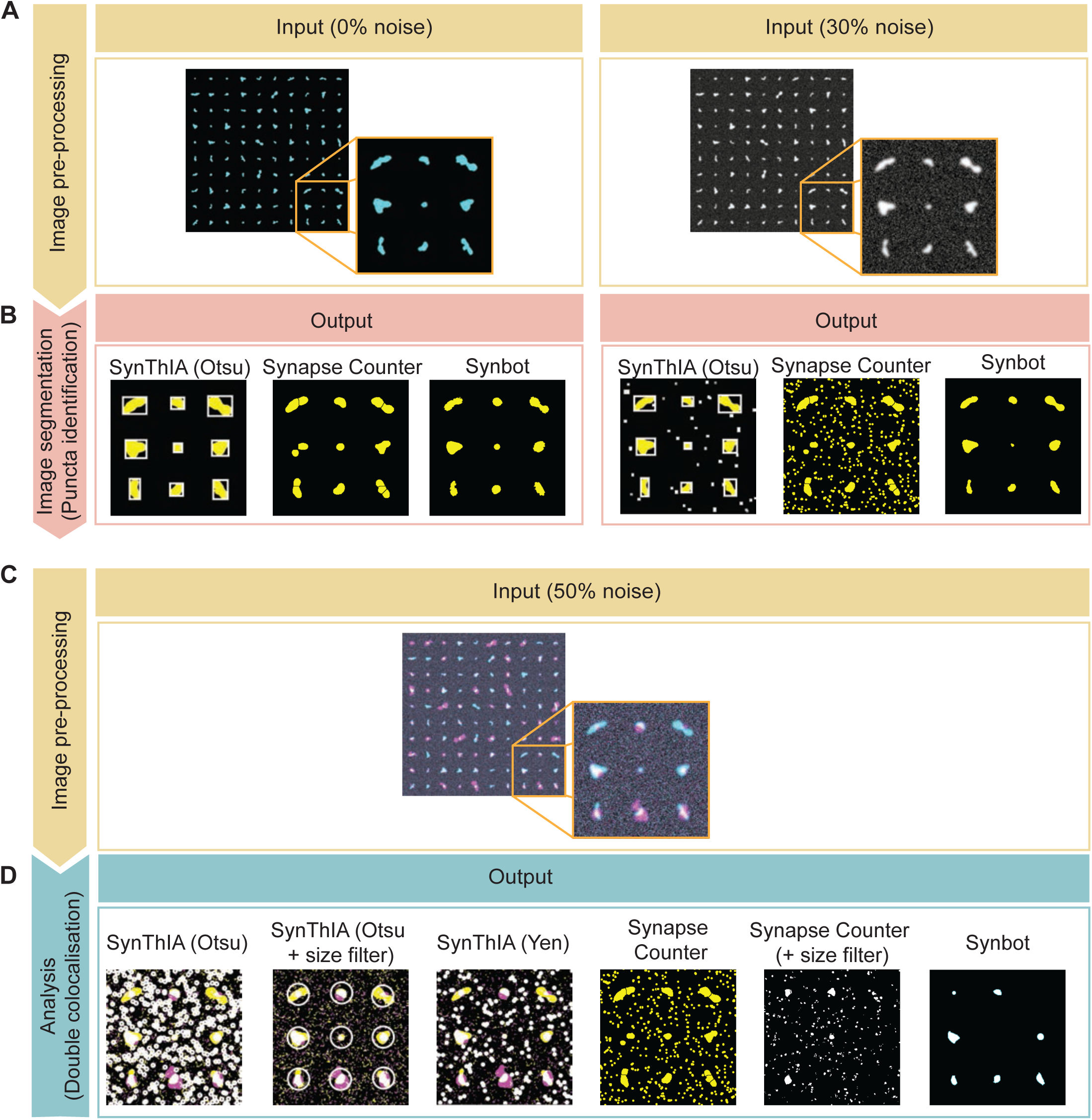
Example of output images using simulated synaptosome images related to Figure 2. (**A**) Schematic of the pipeline on the image pre-processing step with 0% noise (left) and 30% noise (right) (**B**) Example of outputs at the image segmentation from SynThIA (Otsu), Synapse counter, and SynBot with 0% noise (left) and 30% noise (right). Note: White boxes in SynThIA (Otsu) denote identified puncta within the field of view. For Synapse Counter and SynBot, the displayed yellow puncta denote identified puncta within the field of view. (**C**) Schematic of the pipeline on the image pre-processing step with 50% noise. (**D**) Example of outputs of double colocalization identified from SynThIA (Otsu), SynThIA (Otsu) with size filtering, SynThIA (Yen), Synapse Counter, Synapse Counter with size filtering, and SynBot. Note: White circles in SynThIA outputs denote identified puncta within the field of view. For Synapse Counter and Synbot, displayed puncta denote identified puncta within the field of view.

**Figure S3.**
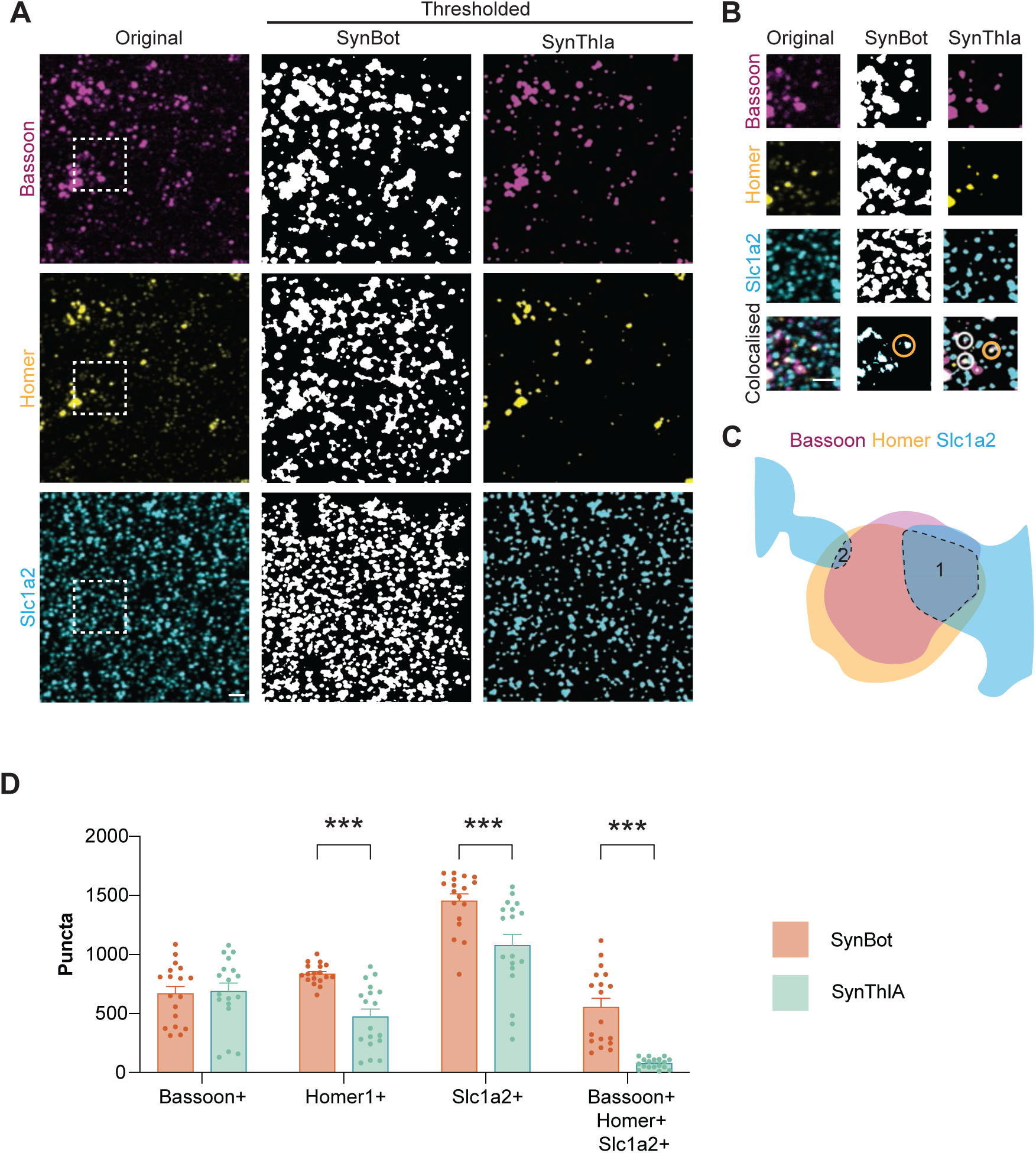
Comparison between SynThIA and SynBot in quantifying isolated synaptosomes related to Figure 3. (**A**) Cortical synaptosomes isolation from Sighted-FVB mice at P56 following immunohistochemistry for Bassoon (magenta), Homer (yellow), and Slc1a2 (cyan) (left), together with thresholded output from SynBot (middle) and SynThIA (right). Dashed white boxes showed the amplified area in (B). (**B**) Higher magnification of cortical synaptosomes isolation from Sighted-FVB from (A) together with the thresholded output from SynBot (middle) and SynThIA (right). Circles showed identified colocalised puncta. The identified colocalised punctum is shown for the yellow circle in (C). (**C**) Schematic drawing of double counting in SynBot. (**D**) Quantification of the total number of Bassoon+, Homer+, Slc1a2+, and triple-colocalised puncta quantified by SynBot (orange) and SynThIA (green). 2-way ANOVA (F*_interaction_* (3, 136) = 5657, *** *p* = 0.0002 (Homer1), *** *p* = 0.0001 (Slc1a2), *** *p* < 0.0001 (triple colocalization), N = 18 wells from 6 mice). The data points within the bar graphs indicate the number of events quantified per image. Scale bar: 5µm,

**Table S1.**
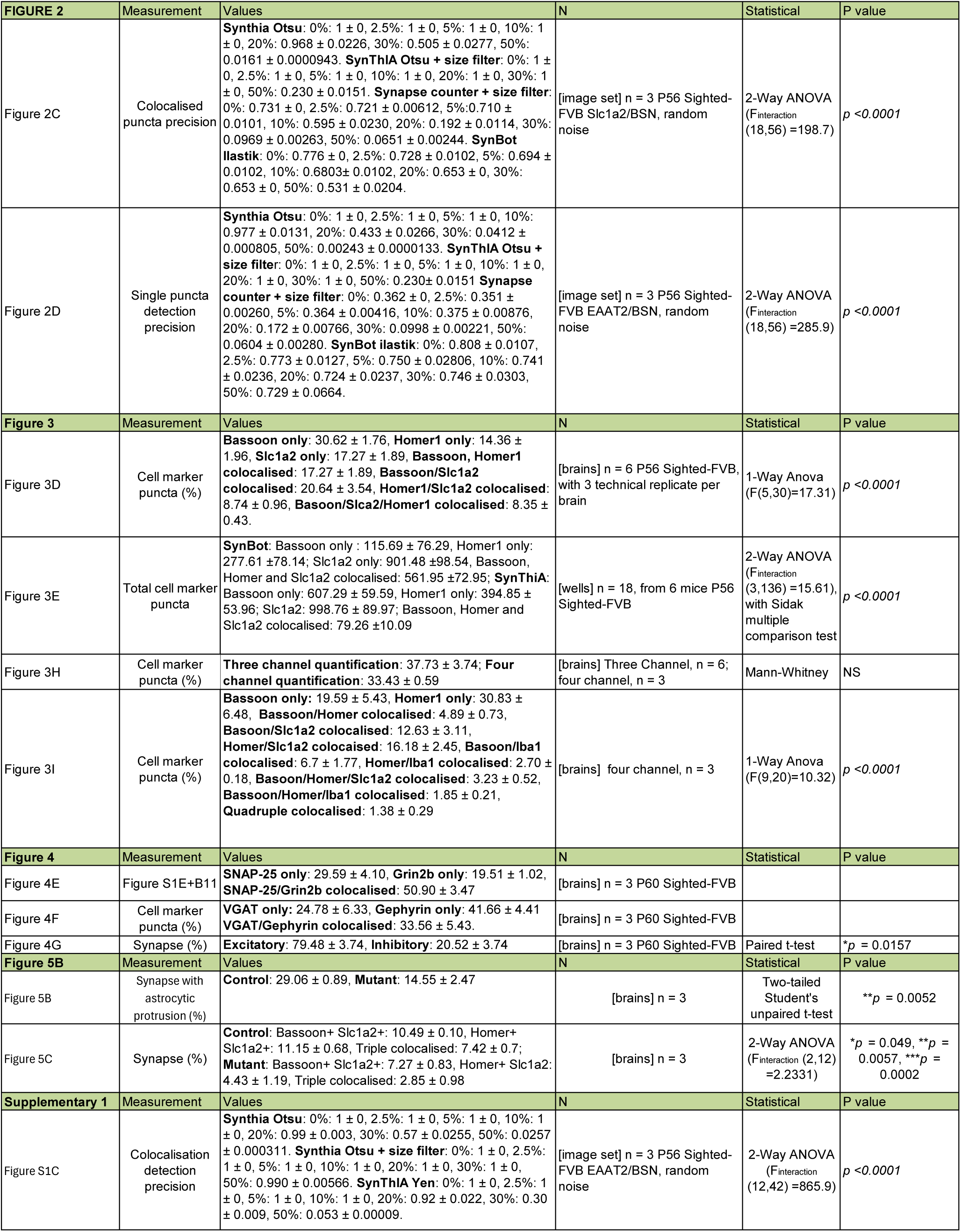
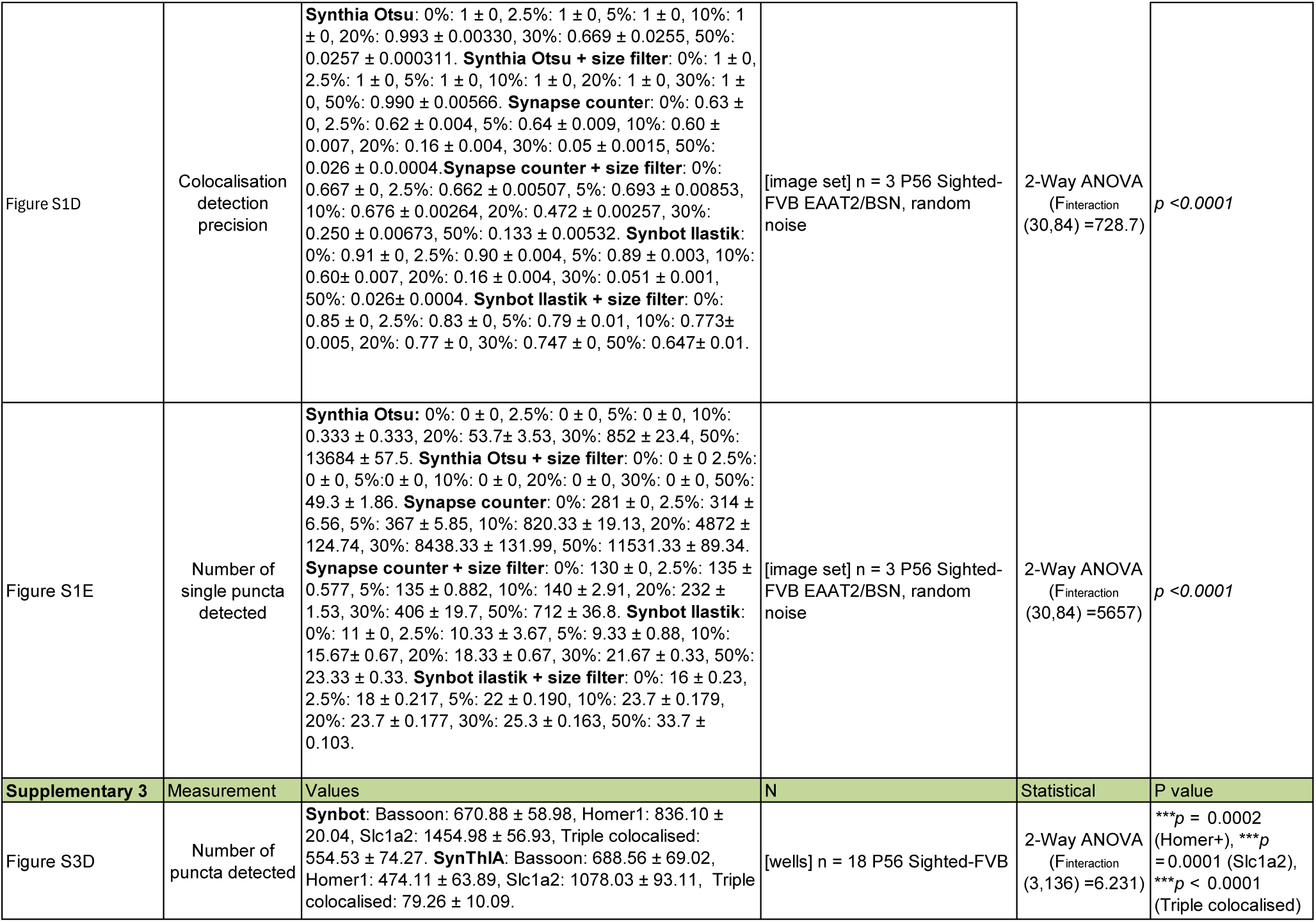
Summary of data and statistical analyses, related to Figures 1-6 and Figures S1-S18.

**Table S2.** Comparison table of key features among Synapse Counter, SynBot and SynThIA.

| Programme name | Synapse Counter | SynBot | SynThIA |
| --- | --- | --- | --- |
| Year of publication | 2017 | 2024 | 2025 |
| Number of channels | 2 | 3 | 4 |
| Availability | https://github.com/SynPuCo/SynapseCounter | https://github.com/Eroglu-Lab/Syn_Bot | https://github.com/FKW-Lab/SynThIA |
| Installation | Preinstalled to FIJI | Install to FIJI through online download. Additional downloads required for Ilastik and SynQuant | Python install required with installation of dependencies |
| Denoising | Modifiable rolling ball + median filter | Preset gaussian blur + rolling ball | Difference of gaussian+ rolling ball or non-local means + median filter |
| Thresholding | Multiple FIJI thresholding options | Ilastik / Synquant / FIJI integrated / manual | Multiple global options / manual / SynQuant algorithm |
| Segmentation accuracy | Dependent on user threshold selection | Dependent on Ilastik training quality | Dependent on threshold selection (further threshold integration possible) |
| Colocalisation methodology | Pixel overlap | Pixel overlap / circular approximation (2 channels only) | Circular approximation with automated threshold |
| Time per image set (based on the 19 image sets processed for s1) | 3 seconds | 81 seconds | 13 seconds |
| Training data | None required (user optimised based on image output) | Training files required for Ilastik thresholding | None required (user optimised based on image output) |
| Experience required | No experience required | Some training required for Ilastik use | Some python experience required for modification |
| Output | Excel file of colocalisation, intensity, single channel counts. Thresholded images + colocalised pixel images | Excel files of single channel counts, excel summary file of colocalisation, images of single channel segmentation + colocalised pixels | Excel file with counts for colocalisation across each channel and single channels + images of each processing and colocalisation step |
| Platform | Java / FIJI plugin | Java / FIJI plugin | Python script |
| Customisability | Limited without further plugin integration | Modifiable Java script – coding knowledge required | Scikit library preinstalled for additional thresholding functions |

## STAR METHODS

### KEY RESOURCES TABLE

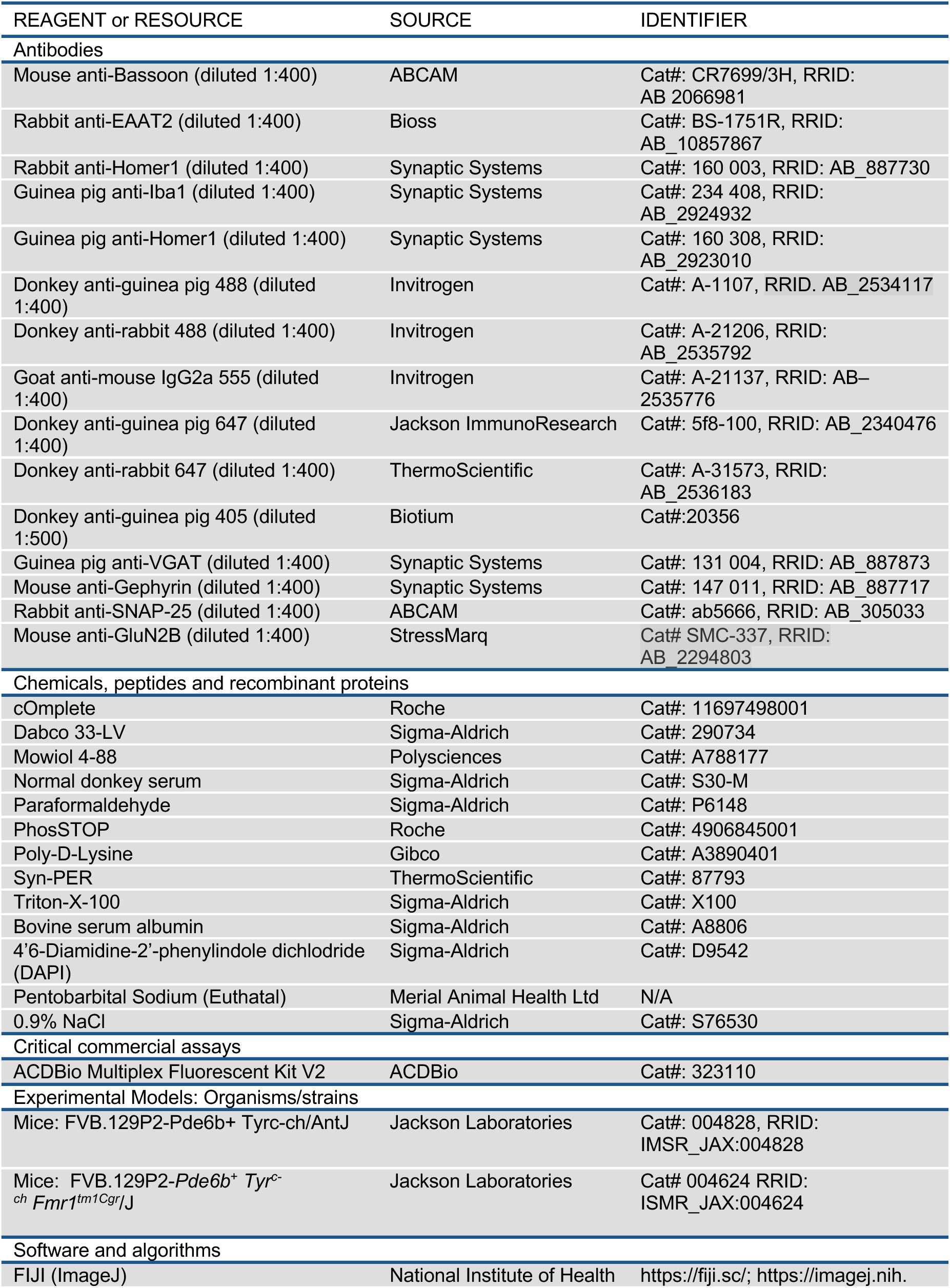
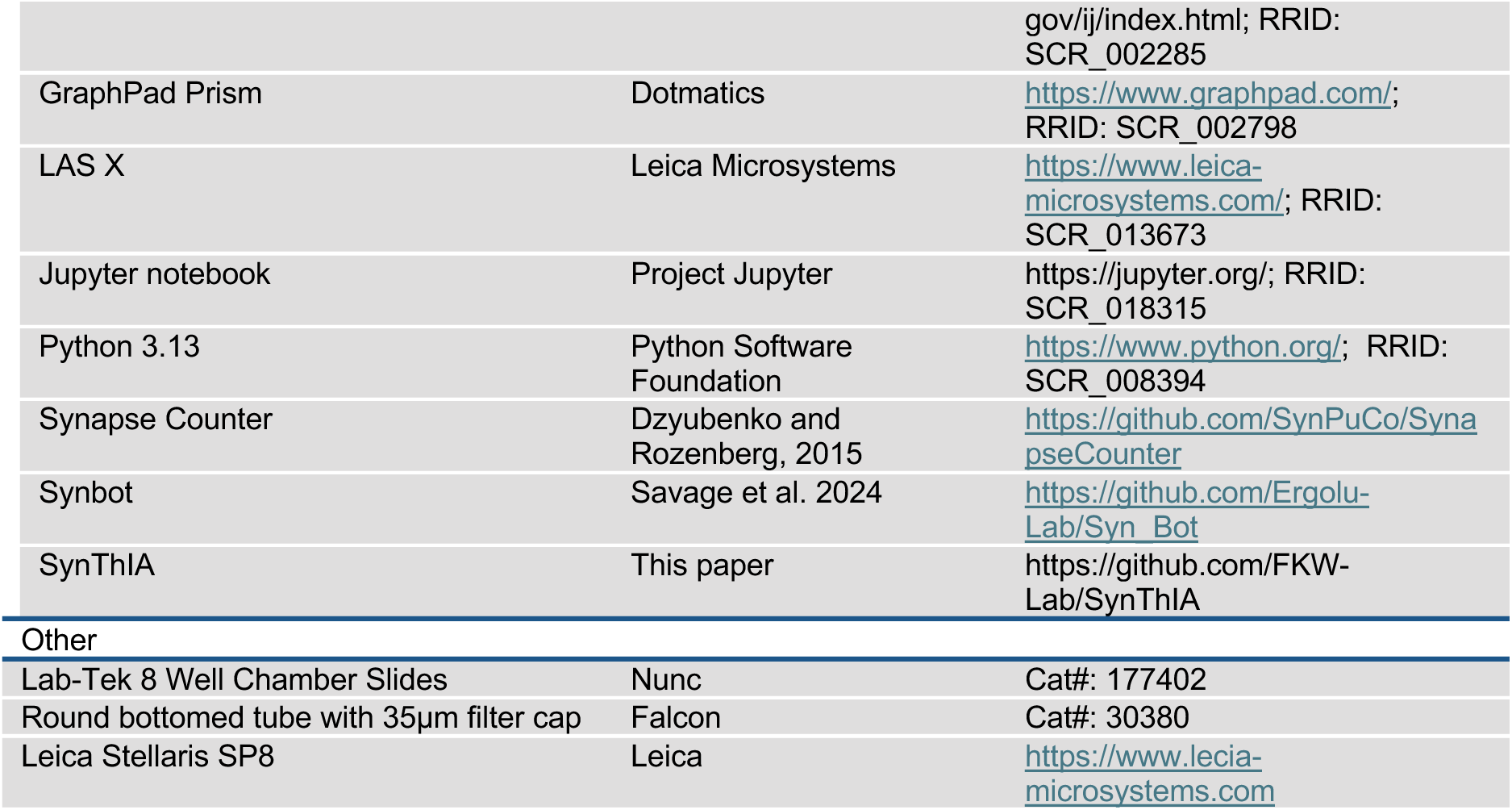

### CONTACT FOR REAGENT AND RESOURCES SHARING

Further information and requests for reagents may be directed to and will be fulfilled by the lead contact, Fong Kuan Wong

### EXPERIMENTAL MODELS

#### Mice

The mouse line Sighted FVB (FVB129P2-Pde6b+ Tyrc-ch/AntJ)^35^ and FMR1 KO (FVB.129P2-Pde6b^+^ Tyr^c-ch^ Fmr1^tm1Cgr^/J)^36^ were used in this study. Animals were housed in groups of up to five littermates and maintained under standard, temperature-controlled, laboratory conditions. Mice were kept on a 12:12 light/dark cycle and received water and food ad libitum. All animal procedures were approved by the ethical committee (University of Manchester) and conducted following European regulations, and Hope Office persona; and project licenses under the UK Animals (Scientific Procedures) 1982 Act.

### METHODS DETAILS

#### Histology

##### Immunohistochemistry

Mice were anaesthetised with an overdose of sodium pentobarbital and transcardially perfused with saline followed by 4% paraformaldehyde (PFA). Brains were post-fixed for 2 h at 4°C. Brains were sectioned using a vibratome (Leica Vibratome VT1000S) at 60µm. Sections were permeabilised using 0.5% Triton-X-100, 10% horse serum and 2% BSA. All primary and secondary antibodies were diluted in PBS containing 10% horse serum and 2% BSA. The following antibodies were used mouse anti-Bassoon (1:400, Abcam), rabbit anti Homer-1 (1:400, Synaptic System) and Rabbit anti-Eaat2 (1:400, Bioss). We used Alexa Fluor-conjugated secondary antibodies (Jackson ImmunoResearch and Invitrogen). Samples were mounted in Mowiol/DABCO.

#### Synaptosome experiments

##### Synaptosome preparation

Cortical synaptosomes from both hemispheres were dissected in ice cold PBS. Cortices were immediately homogenised in SynPER Synaptic Protein Extraction Reagent (ThermoScientific) with added cOmplete protease inhibitors (Roche) and phosSTOP phosphatase inhibitors (Roche). Synaptosomes were prepared according to the manufacturer’s instructions and processed for further applications.

##### Synaptosome plating and immunofluorescence

Cortical synaptosomes were incubated for 60 min at 37°C with gentle shaking on Nunc Lab-Tek 8-well chamber slides (ThermoScientific) coated with 35% poly-D-lysine (Gibco). Synaptosomes were fixed with 4% PFA for 15 min at room temperature and washed with PBS. Synaptosomes were permeabilised in 0.5% Triton-X in PBS for 10 min at room temperature. Samples were then blocked in 4% horse serum in PBS. Samples were then incubated overnight at 4°C with primary antibodies. The following primary antibodies were used: mouse anti-Bassoon (Abcam), rabbit anti-Homer1 (Synaptic System), guinea pig anti-Homer-1 (Synaptic System), rabbit anti-Eaat2 (Bioss), guinea pig anti-Iba1 (Synaptic System). The following day, samples were washed in PBS and incubated with secondary antibodies for 1 h. We used Alexa Fluor-conjugated secondary antibodies (Jackson ImmunoResearch and Invitrogen). Samples were then washed in PBS and mounted with Mowiol/DABCO.

#### Imaging and analysis

##### Image acquisition

Images used for analysis were obtained on the Stellaris 8 confocal microscope (Leica) using the LAS X software. Samples from the same experiment were image and analysed in parallel, using the same laser powers and detection filter settings with similar photo-multiplied gain. Imaging of synaptic markers and in situ hybridisation was performed at 8-bit depth, with 40x objective and 5.5 digital zoom at 200Hz acquisition speed with a line accuracy of 4.

##### Creation of simulated synapse images

Simulated images were produced by taking 30 random puncta from images obtained from plated cortical synaptosomes stained for Bassoon and Eaat2. These puncta were denoised and processed using Otsu thresholding. The 30 reference puncta were then pasted on 10 x 10 grids and duplicated until the grids were filled. Duplicated puncta were randomly rotated to avoid repetition in colocalisation across channels. Two sets of simulated synapse images were created – 1) images containing 100 overlapping puncta and 2) images where 30 puncta were randomly deleted from the grid in each channel. In this set, there are 49 overlapping puncta, 21 single puncta in each channel and 9 blank spaces. All images then had a gaussian blur of sigma = 3 applied to conceal the edges of the puncta. The blurred images then had salt noise randomly added to 2.5, 5, 10, 20, 30 and 50% of the pixels in the image. The intensity of this salt noise was also randomised to any possible greyscale value. Three distinct sets of images were produced.

##### Simulated synapse quantification with SynThIA

SynThIA was installed in a virtual environment and run in JupyterLab as per the installation instructions. Images were analysed using the 2 channel colocalisation function with the images saved as individual channels using the naming convention outlined in the instructions. Thresholding was set as “no background” and either “Otsu” or “Yen”. Where size filtering was combined with Otsu thresholding, the size filtering box was selected and a size filter of 50 – 3000 pixels applied. This was calculated through approximate measurement using the ImageJ measure function.

##### Simulated synapse quantification with Synapse Counter

Individual images for each channel were combined into hyperstacks using Fiji and loaded to Synapse Counter^12^. The settings were kept to default, with a rolling ball radius of 10, maximum filter radius of 2 and Otsu threshold adjustment. For quantification without size filtering, filter size was set at 0 -20 000 pixels. For quantification with size filtering, filter size was the same as used with SynThIA (50 – 3000 pixels).

##### Simulated synapse quantification with SynBot

2 additional sets of 14 images containing two channels were generated to form a training set for the Ilastik thresholding option within SynBot. Ilastik was then trained on the images as per instructions and loaded to SynBot to threshold the images.

##### Synaptosome quantifications

Synaptosome images were quantified by SynThIA directly from LIF files with the image processing set as no background, Otsu thresholding and a 5-200 size filter, in line with typical size filtering used for synaptosome images. The image were quantified using Synapse Counter with the default synapse counter settings, including the 5-200 size filter. For SynBot, images were quantified using an ilastik profile trained on the images themselves without any size filtering.

##### Tissue image quantification

Tissue images were quantified by SynThIA directly from .LIF files using the 3-channel function, with ‘background’ selected and a rolling ball radius of 10 for the Bassoon channel and 25 for the remaining channels. Otsu thresholding and a 5< size filter were used. The LIF file was split into hyperstacks of each set of 3 channels for processing with Synapse Counter and SynBot. Synapse Counter with default settings were used. SynBot was used with default settings and ilastik thresholding trained on five example images.

#### Statistical Analysis

All statistical analysis were performed using Prism 10 (GraphPad Software). Samples were tested for normality using the Shapiro-Wilk normality test. Paired comparisons were analysed using two-tailed unpaired Student’s *t*-test (normally distributed) and Mann-Whitney test (for not normally distributed). Multiple comparisons with a single variable were analysed using one-way ANOVA with post-hoc Tukey. Statistical significance was considered at p-values ≤ 0.05. The number of animals for each experiment is described in each figure legend.

